# Influence of substrate stiffness on barrier function in an iPSC-derived in vitro blood-brain barrier model

**DOI:** 10.1101/2021.03.31.437924

**Authors:** Allison M. Bosworth, Hyosung Kim, Kristin P. O’Grady, Isabella Richter, Lynn Lee, Brian J. O’Grady, Ethan S. Lippmann

## Abstract

**Introduction:** Vascular endothelial cells respond to a variety of biophysical cues such as shear stress and substrate stiffness. In peripheral vasculature, extracellular matrix (ECM) stiffening alters barrier function, leading to increased vascular permeability in atherosclerosis and pulmonary edema. The effect of ECM stiffness on blood-brain barrier (BBB) endothelial cells, however, has not been explored. To investigate this topic, we incorporated hydrogel substrates into an *in vitro* model of the human BBB.

**Methods:** Induced pluripotent stem cells were differentiated to brain microvascular endothelial-like (BMEC-like) cells and cultured on hydrogel substrates of varying stiffness. Cellular changes were measured by imaging, functional assays such as transendothelial electrical resistance (TEER) and p-glycoprotein efflux activity, and bulk transcriptome readouts.

**Results:** The magnitude and longevity of TEER in iPSC-derived BMEC-like cells is enhanced on compliant substrates. Quantitative imaging shows that BMEC-like cells form fewer intracellular actin stress fibers on substrates of intermediate stiffness (20 kPa relative to 1 kPa and 150 kPa). Chemical induction of actin polymerization leads to a rapid decline in TEER, agreeing with imaging readouts. P-glycoprotein activity is unaffected by substrate stiffness. Modest differences in RNA expression corresponding to specific signaling pathways were observed as a function of substrate stiffness.

**Conclusions:** iPSC-derived BMEC-like cells exhibit differences in passive but not active barrier function in response to substrate stiffness. These findings may provide insight into BBB dysfunction during neurodegeneration, as well as aid in the optimization of more complex threedimensional neurovascular models utilizing compliant hydrogels.

## INTRODUCTION

Endothelial cells (ECs) line blood vessels and perform vital organ-specific functions. Within the blood vessel architecture, ECs sit on an intimal basement membrane which is surrounded by smooth muscle cells or pericytes depending on the location along the vascular tree. Due to their barrier properties, ECs play a key role in shuttling substances between the parenchyma and the bloodstream. ECs are known to respond to a variety of extracellular mechanical cues^5^, including shear flow, bifurcations, topography, curvature, matrix composition, and stiffness^8,14,32^. These properties can impart a positive or negative effect on EC barrier function, and the sensitivity of ECs to mechanical cues can impact a variety of disease states^8^.

Stiffness, in particular, has been shown to increase permeability of endothelial barriers and limit contraction of larger vessels. Across all vessel sizes, there are reports of ubiquitous changes in EC function as the underlying substrate becomes stiffer. For example, *in vitro* experiments have been used to demonstrate that, when placed on stiffer substrates, aortic endothelial cells exhibit increased permeability and leukocyte transmigration^13^, and lung microvascular endothelial cells exhibit decreased transendothelial electrical resistance (TEER) and discontinuous junctions^22^. Pulse wave velocity measurements of human vessels have shown that macroscale vessel stiffness increases in aging and atherosclerosis^6^, and these changes disrupt endothelial barriers^35^. Some of the mechanisms that transduce mechanical cues into barrier modifications have been illustrated as well, for example that cellular cytoskeleton and contractility are altered on stiffer matrices and negatively affect endothelial permeability^1,13^. However, the connections between stiffness and endothelial barrier function have thus far only been assessed for peripheral endothelial cells and not those with more stringent barrier properties, such as specialized brain microvascular endothelial cells (BMECs) that form the blood-brain barrier (BBB). BMECs are distinguished from peripheral ECs by the presence of complex tight junctions, higher mitochondrial content, lack of fenestrations, and very little pinocytosis^10^, which are critical features that help maintain neural function and brain homeostasis^31^. BMECs exhibit very high TEER, highlighting their ability to limit molecular transport along paracellular pathways. Additionally, BMECs express a variety of membrane transporters to actively shuttle molecules into and out of the brain along transcellular pathways. Altogether, these properties allow the BBB to strictly regulate the transport of compounds between the bloodstream and the brain. At present, it has not been assessed whether BMECs and their specialized BBB properties are influenced by underlying substrate stiffness.

BMECs have historically been difficult to study *in vitro* because they lose BBB characteristics after removal from the *in vivo* microenvironment^29^. This challenge has been partially circumvented in recent years by the advent of human pluripotent stem cell technology, which has permitted the generation of endothelial-like cells with robust BBB properties^20,21^. Previous work from our group demonstrated that these human induced pluripotent stem cell (iPSC)-derived BMEC-like cells respond to shear stress when cultured as a perfusable macrovessel structure. When cultured under constant shear, iPSC-derived BMEC-like cells were 10-100 times less permeable to 3 kDa dextran relative to those cultured under static conditions.^7^ Different perfusion rates also yielded changes to dextran permeability and induced angiogenic-like sprouting into the surrounding hydrogel, which indicated that iPSC-derived BMEC-like cells are sensitive to mechanical cues. However, the response of iPSC-derived BMEC-like cells to the stiffness of the underlying substrate was not explicitly tested. Here, we sought to further characterize how iPSC-derived BMEC-like cells respond to stiffness cues using hydrogels, functional assays, and molecular biology tools. We determined that, in agreement with general mechanobiology trends, iPSC-derived BMEC-like cells develop tighter barriers on hydrogels with intermediate stiffness that match healthy vessel intima. These cells reorganize actin into intracellular stress fibers when substrate conditions are too soft or too stiff, and small molecule-mediated induction of actin polymerization yielded a rapid reduction in TEER, suggesting links between actin structure and passive BBB function. Finally, RNA sequencing was used to provide further insight into how iPSC-derived BMEC-like cells respond to culture on substrates of varying stiffness. We suggest that our findings will be useful towards fabricating more complex 3D BBB models that involve hydrogels and may be interesting for studies of vascular dysfunction in neurodegeneration.

## METHODS

### Polyacrylamide Hydrogel Preparation

For coverslip-bound hydrogels, activated coverslips and polyacrylamide (PA) hydrogels were prepared as previously reported^4^. 15 mm circular glass coverslips (Carolina 633011) were exposed to oxygen plasma for 6 minutes, incubated with a 0.1% polyethyleneimine (Sigma P3143) solution for 10 minutes, washed three times in deionized water for 5 minutes each, and incubated with 0.5% glutaraldehyde (Sigma G7776) for 30 minutes. Three additional 5-minute washes were performed, and activated coverslips were dried overnight in fume hood.

Stock 40% acrylamide (BioRad 1610140) and 2% bis-acrylamide (BioRad 1610143) solutions were diluted in ultrapure water at varying concentrations to yield hydrogels of varying stiffness. Final concentrations of 3% A and 0.1% B, 15% A and 0.1% B, 15% A and 1.2% B were used to fabricate hydrogels at approximately 1 kPa, 20 kPa, and 150 kPa, respectively. To fabricate the gels, these PA solutions were degassed for 1 hour, then TEMED (BioRad 1610801) and ammonium persulfate (APS) (BioRad 1610700) were added to initiate polymerization. For each milliliter of PA solution, 1 μl of TEMED and 5 μl of a freshly reconstituted 10% APS solution were added. 35 μl of this solution was immediately sandwiched between a Rain-X-coated microscope slide and the activated coverslip. Polymerization occurred for 20 minutes before gels were unmolded to reveal a coverslip-bound hydrogel of 0.2 mm thickness.

Hydrogels were incubated in a 50 mM HEPES (Sigma 391340) buffer of pH 8 for at least 30 minutes on a rocking plate at room temperature before being coated with a sulfo-SANPAH (SS) (CovaChem 13414-5×5) linker. SS was reconstituted in aforementioned HEPES buffer at 0.5 mg/ml and immediately pipetted onto the hydrogel surface. PA hydrogels were exposed to 110 mJ/cm^2^ of UV light (UVP CL-1000 UV Crosslinker) for 10 minutes. Once this exposure was completed, fresh SS was added to the hydrogels, and a second UV exposure was performed. Then, PA hydrogels were washed twice in HEPES buffer for 15 minutes on a rocking platform to remove excess, unbound SS. Following the washes, the PA hydrogels were incubated with extracellular matrix protein (ECM) solutions overnight at 4°C. To prepare the protein solution, collagen type IV (Sigma C5533) and fibronectin (Sigma 1141) were diluted in sterile HEPES buffer to final concentrations of 0.4 and 0.1 mg/ml, respectively.

After overnight incubation, hydrogels were submerged in 5% PenStrep for 4 hours at room temperature in preparation for cell seeding. After 4 hours, hydrogels were rinsed 3 times with sterile, ultrapure water, then incubated with DMEM/F12 for 30-60 minutes at 37°C prior to cell seeding.

### Gelatin Hydrogel Fabrication

Gelatin from porcine skin (Sigma G1890) was reconstituted in water at 10% (w/v), then further diluted to produce 7.5%, 5%, and 2.5% gelatin solutions. 10 mL of each solution was mixed with 1 mL of a 10% (w/v) solution of microbial transglutaminase (Modernist Pantry 120-350) in water. This precursor solution was thoroughly mixed by pipetting. To polymerize hydrogels in Transwell filters, 350 uL of precursor solution was pipetted onto the top of the filter. To produce hydrogels bound to coverslips, 130 uL of precursor solution was sandwiched between Rain-X-coated microscope slides and glutaraldehyde-activated coverslips. For both Transwell and coverslip formats, precursor solutions were incubated for 1 hour at 37°C to crosslink. Hydrogels on Transwells were incubated with ECM solution overnight for adsorption in preparation for cell seeding. ECM solution was prepared at same concentrations as mentioned above, but proteins were diluted in sterile water instead of HEPES buffer. Coverslip hydrogels were not coated with ECM.

### Quantifying Stiffness of Gels by Atomic Force Microscopy

Stiffness of hydrogels was quantified using a Bruker Dimension Icon Atomic Force Microscope in the Vanderbilt Institute for Nanoscale Science and Engineering. Both sulfo-SANPAH PA hydrogels and gelatin hydrogels were prepared as coverslip-bound gels for atomic force microscopy (AFM) measurements. Coverslips were glued to microscope slides and measurements were performed in fluid. PA hydrogels were measured using Bruker precalibrated PFQNM-LC-A probes with a nominal spring constant of 0.1 N/m and a tip radius of 70 nm, and gelatin hydrogels were measured using Novascan pre-calibrated PT.PS.SN.4.5.CAL probes with a nominal spring constant of 0.01 N/m and a tip radius of 2.25 μm. Three distinct 5×5 micron areas were measured across three hydrogels for each condition, totaling 9 captures per condition. The resulting force curves for each condition were baseline corrected, and Young’s Modulus was calculated according the Hertzian model. Values for each condition were averaged to produce the reported stiffness values found in Figures 1 and 2.

**Figure 1.**
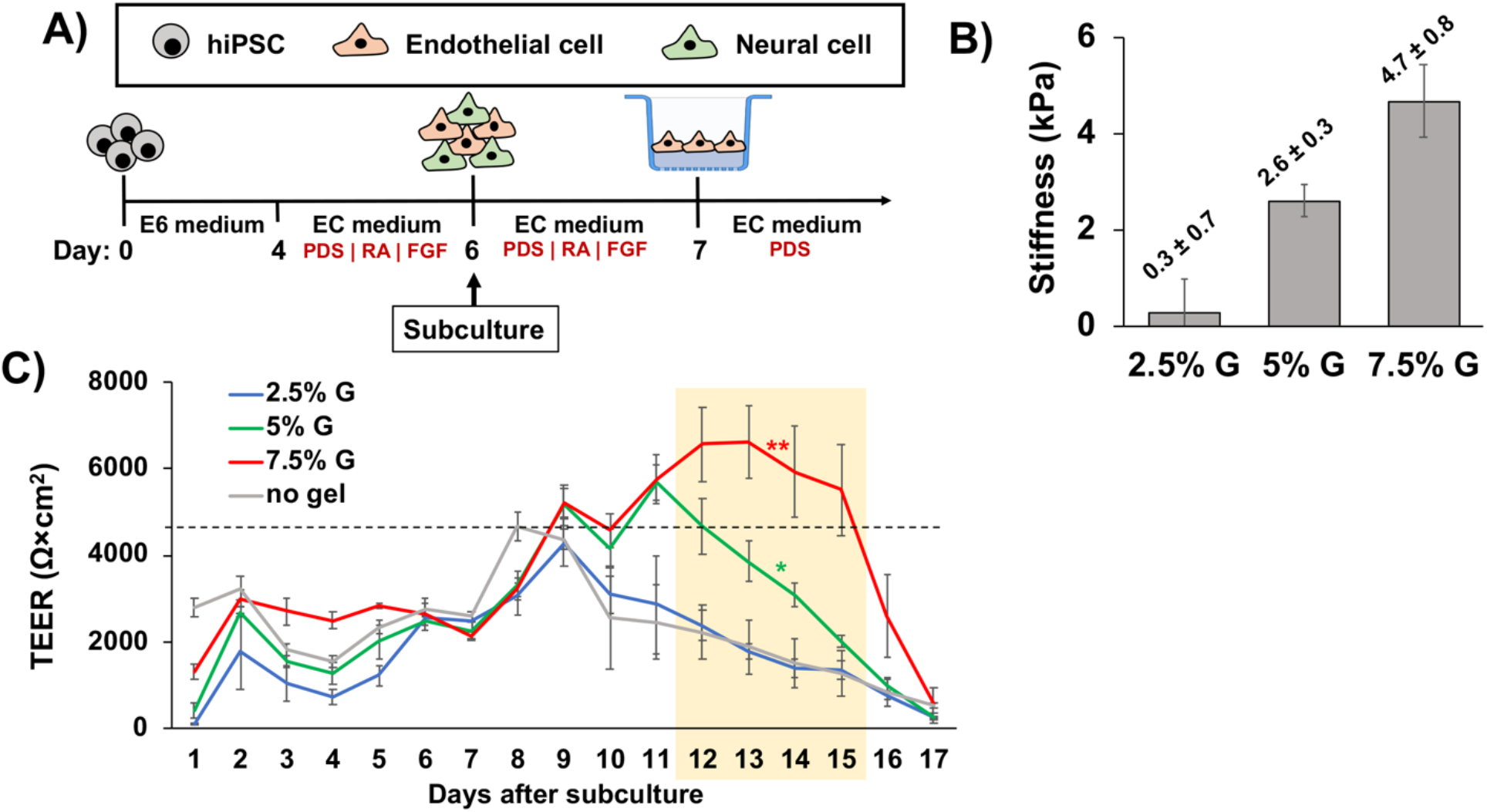
iPSC-derived BMEC-like cells exhibit optimal passive barrier properties on compliant substrates. (A) Differentiation and seeding strategy on gelatin hydrogels. (B) Stiffness of the gelatin hydrogels of varying weight percent. AFM data are presented as mean ± standard deviation from 9 total measurements (3 separate hydrogels, 3 measurements per hydrogel). (C) Representative TEER plot for iPSC-derived BMEC-like cells cultured on gelatin hydrogels. 3 filters were prepared for each condition, and each filter was measured in triplicate. Values for each day represent mean ± standard deviation from these 9 measurements. The dotted line indicates maximum average TEER achieved by the no hydrogel control. A one-way ANOVA with a Tukey post hoc test was performed for days 12-15 (yellow box), which showed that the 7.5% and 5% gelatin hydrogels had significantly elevated TEER relative to the 2.5% gelatin hydrogel and no hydrogel control on each day (\**p*<*0.05*, \*\**p*<*0.01*). These trends were confirmed across an additional biological replicate.

**Figure 2.**
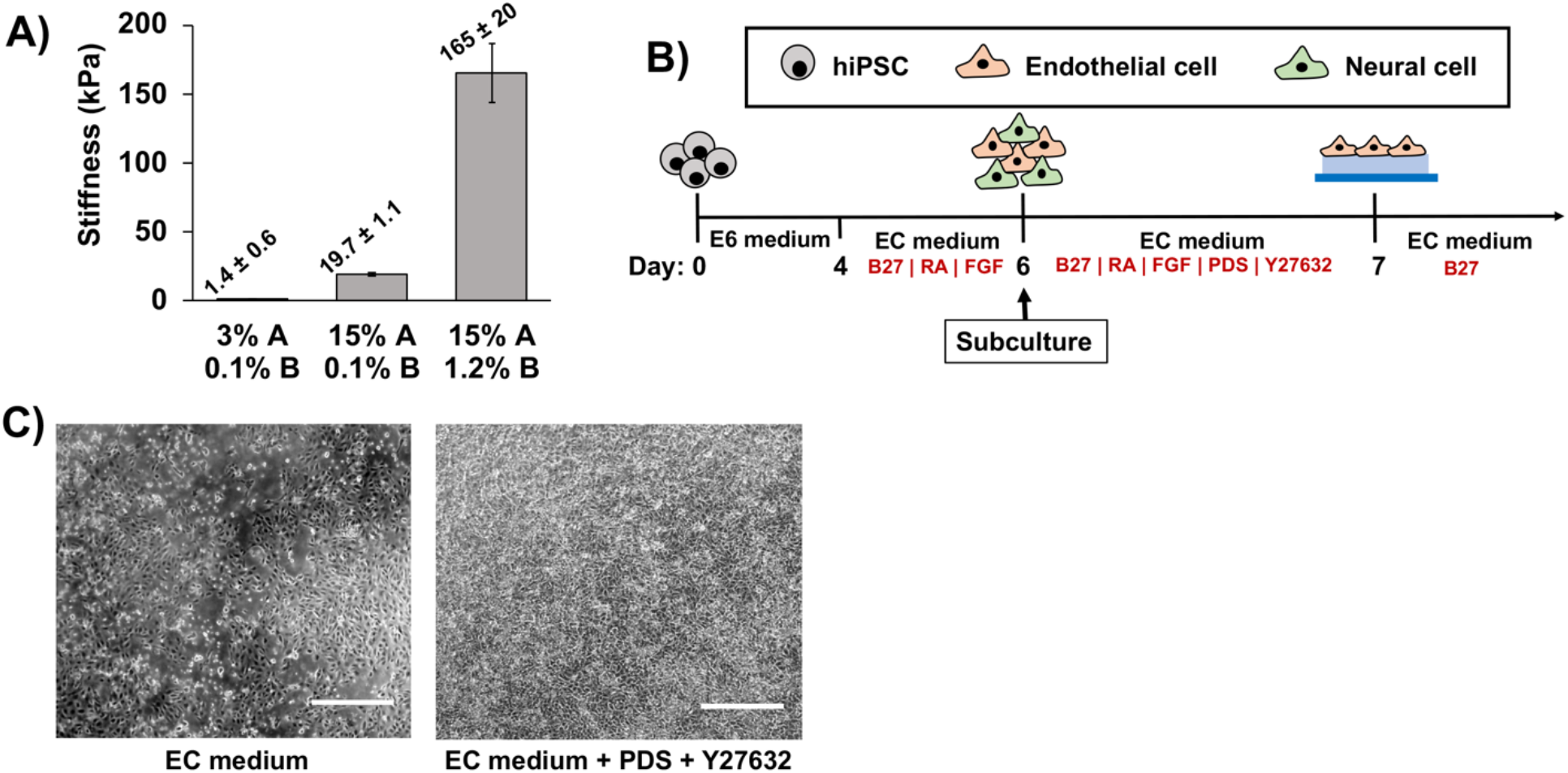
Strategy for seeding iPSC-derived BMEC-like cells as confluent monolayers on PA hydrogel substrates. (A) Stiffness of the PA hydrogels synthesized with varying acrylamide (“A”) and bisacrylamide (“B”) ratios. AFM data are presented as mean ± standard deviation from 9 total measurements (3 separate hydrogels, 3 measurements per hydrogel). (B) Differentiation and seeding strategy on PA hydrogels. (C) Brightfield images showing that iPSC-derived BMEC-like cells sporadically adhere to PA hydrogel surfaces in base EC medium but form confluent monolayers when base EC medium is supplemented with PDS and Y27632 for 24 hours. Scale bars are 250 μm.

### Differentiation of iPSCs to BMEC-like cells

iPSCs were maintained and differentiated as previously described^11,26^. For all experiments, to initiate differentiation, CC3 iPSCs were seeded in E8 medium containing 10 μM Y27632 (Fisher Scientific 12-541-0) at 12,500 cells/cm^2^. Then, 24 hours after seeding, cells were switched to E6 medium and daily media changes were performed for four days.

From this point, different routes were taken depending on the experiment and hydrogel substrate. For experiments on gelatin hydrogels, we started with an older differentiation procedure that utilized serum-containing medium^11^. Briefly, on day 4 of differentiation, media was changed to the following formulation of EC medium: human endothelial serum-free medium (hESFM, Thermo Fisher 1111044) supplemented with 1% platelet-poor plasma-derived serum (PDS, Fisher AAJ64483AE), 10 μM retinoic acid (RA, Sigma R2625), and 20 ng/mL basic fibroblast growth factor (bFGF, Peprotech 100-18B). No media change was performed on day 5. On day 6, cells were dissociated in Accutase for 15-20 minutes (or until approximately 70% of cells had visibly dissociated), centrifuged, and replated in the same EC medium onto gelatin hydrogels cast in Transwell filters, as described above. Cells from 3 wells of a 6-well plate were split across 9 Transwell filters. 24 hours later, media was aspirated and changed to fresh EC medium containing PDS but without RA or bFGF. No further media changes were performed.

For experiments on PA hydrogels and standard Transwell filters, we utilized newer variations of our protocol that remove serum from the differentiation procedure^26,27^. Here, on day 4 of differentiation, media was switched to updated EC medium: Neurobasal (Thermo Fisher 21103049) with 0.25% GlutaMAX (Thermo Fisher 35050061), 0.5% B27 supplement (Thermo Fisher 17504044), 10 μM RA, and 20 ng/mL bFGF. No media change was performed on day 5. On day 6, cells were dissociated in Accutase for 15-20 minutes (or until approximately 70% of cells had visibly dissociated), centrifuged, and replated onto polyacrylamide hydrogels or standard Transwell filters. If seeding onto hydrogels, cells were plated in the aforementioned EC medium containing 0.1% PDS and 1 μM Y27632 (Tocris 1254) to facilitate adhesion. If seeding onto Transwell filters coated with 0.4 mg/ml collagen IV and 0.1 mg/ml fibronectin, cells were plated in the same EC medium utilized at day 4. Cells from 2 wells of a 6-well plate were split across 3 hydrogels or 6 Transwell filters. 24 hours later, media was aspirated and changed to fresh EC medium containing B27 but without RA, bFGF, PDS, and Y27632. No further media changes were performed.

### TEER measurements

For experiments on gelatin hydrogel substrates, TEER was measured using an EVOM2 voltohmeter with EndOhm cup chamber (World Precision Instruments ENDOHM-12g) filled with pre-warmed hESFM medium. Each day, height of the upper electrode was set according to an empty hydrogel Transwell containing no cells. Resistance values were recorded for the empty hydrogel Transwells and for the sample conditions. For experiments on cells cultured directly on Transwell filters, TEER was measured using an EVOM2 voltohmeter with STX3 chopstick electrodes (World Precision Instruments). 0.13 μM jasplakinolide (jasplak; Thermo Fisher J7473) was added directly to the cell culture medium, and equivalent volumes of DMSO were added as the vehicle control. Resistance values were recorded from an empty Transwell containing no cells and for the sample conditions. For all experiments, empty measurements were subtracted from sample measurements, and the resulting values were multiplied by 1.1 cm^2^ surface area of the filter.

### P-glycoprotein Efflux Assay

Efflux assays were performed on BMEC-like cells 8 days after seeding onto hydrogels. 10 μM Cyclosporin A (Fisher Scientific 11-011-00) was added to the cell culture media and incubated for 1 hour at 37°C. Cells were then incubated with 10 μM Rhodamine 123 (Thermo Fisher R302) and 10 μM Cyclosporin A in fresh EC medium for 1 hour at 37°C. Cells were then rinsed with PBS, scraped off the hydrogel surface and transferred to a microfuge tube containing TryplE (Thermo Fisher 12604013). Triplicate gels of each condition were pooled, and triturations were performed until no visible cell clumps remained. Cells were re-centrifuged and resuspended in flow buffer (PBS containing 5% donkey serum). Flow cytometry was performed using a Guava easy-Cyte benchtop flow cytometer. Events were plotted as forward scatter versus green fluorescence, and rhodamine-positive cells were identified against BMEC-like cells that received no rhodamine or cyclosporin.

### Immunofluorescence

Cells were rinsed twice with PBS and fixed for 10 minutes in 4% paraformaldehyde. Then, cells were washed three times in PBS containing 0.3% Triton-X, referred to as PBS-T, for 5 minutes each. After washes, cells were blocked in PBS-T containing 5% donkey serum at room temperature for at least 1 hour. Cells were then incubated overnight at 4°C with an occludin primary antibody (Thermo Fisher 33-1500) at 1:200 in PBS containing 5% donkey serum. On the following day, cells were washed five times in PBS-T at 8 minutes each. A donkey anti-mouse Alexa Fluor 647 secondary antibody (Thermo Fisher A31571) was diluted at 1:200 in PBS containing 5% donkey serum and applied to cells for a 1-2 hour incubation at room temperature. Cells were then incubated with Phalloidin-488 (Thermo Fisher A12379) at 1:1000 for 30 minutes and Hoechst (Thermo Fisher H1399) at 1 μg/mL for 10 minutes. Four more washes in PBS-T were performed at 8 minutes each, and cells were mounted onto microscope slides for imaging.

### Image Processing

For qualitative assessment of cell confluence, brightfield images were acquired using an EVOS XL Core Imaging System. For quantification of actin stress fibers and tight junction integrity, phalloidin and occludin were imaged using a Zeiss LSM 880 confocal microscope in the Vanderbilt Cell Imaging Shared Resource Core Facility. Actin stress fibers were manually counted by a blinded observer. Occludin labeling was overlaid during counting to ensure that only intracellular fibers were counted.

For quantification of tight junction width and tortuosity, image processing was performed in Matlab (MathWorks, Inc.). First, fluorescence images of occludin were filtered with a twodimensional Hessian filtering algorithm to enhance contrast using a modification of methods described previously^19,28^. In the filtering step, junctions oriented in any direction were detected using the eigenvalues of the image, and detection was performed over multiple scales. The raw image was then scaled using the maximum filter response for each pixel, and the resulting image was thresholded to produce a binary mask of the tight junctions. The binary mask was skeletonized to find the centerline pixels of all junctions, and the junction width at each centerline pixel was computed. To estimate tortuosity of the junctions, the straight-line distance was computed between branch points in the skeletonized image, and the actual total length of the junctions was divided by the total straight-line distance (ratios closer to 1 indicate junctions are less tortuous).

### RNA sequencing

On day 8 after purification, BMEC-like cells were detached from PA hydrogel surfaces with a 10-minute Accutase incubation at 37°C. Cells from 3 gels of each stiffness condition were pooled to create one biological replicate. Cells were then rinsed with PBS and incubated in Trizol (Thermo Fisher 15596026) for 10 minutes at room temperature. RNA was extracted using a Direct-zol RNA miniprep kit (Zymo R2050) according to the manufacturer’s instruction. This process was repeated two additional times from independent iPSC differentiations (biological n=3).

RNA samples were submitted to the Vanderbilt Technologies for Advanced Genomics (VANTAGE) facility for sequencing using an Illumina NovaSeq6000. Raw sequencing reads were obtained for the 9 paired-end samples. FASTQ reads were mapped to human genome (hg19) by HISAT2 (version 2.1.0). Gene expression levels were estimated through String Tie. EdgeR (version 3.30.3) was used to measure differential gene expression, and genes with p<0.05 were utilized for Gene ontology (GO) enrichment analysis with the WEB-based GEne SeT AnaLysis (WebGestalt) Toolkit^36^. Results were annotated with functional database IDs and false-discovery rates for each signaling pathway.

## RESULTS AND DISCUSSION

### iPSC-derived BMEC-like cells exhibit improved passive barrier function on compliant gelatin hydrogels

To initially examine the influence of stiffness on passive barrier function, we crosslinked gelatin hydrogels in Transwell filters and measured TEER in iPSC-derived BMEC-like cells. We utilized an older variant of our differentiation protocols that contained serum throughout the differentiation process and after purification of the iPSC-derived BMEC-like cells^11^ (Figure 1A). We manipulated the gelatin concentration (2.5, 5, and 7.5 weight percent), which resulted in hydrogel stiffnesses of 0.3, 2.5, and 5 kPa respectively (Figure 1B). Intriguingly, while cells on 2.5% gelatin hydrogels exhibited either similar or lower TEER maxima than cells on control Transwell filters, cells cultured on 5% and 7.5% gelatin exhibited significantly higher TEER maxima, as well as enhanced barrier longevity (Figure 1C). In particular, iPSC-derived BMEC-like cells cultured on 7.5% gelatin hydrogels reach a significantly higher TEER maximum with respect to the softer substrates and the no hydrogel control. These results suggest that passive barrier function of iPSC-derived BMEC-like cells is sensitive to the stiffness of the underlying substrates.

### Optimized seeding of iPSC-derived BMEC-like cells on polyacrylamide hydrogel substrates

The stiffness of ECM-based hydrogels can be tuned by modifying protein concentration, but this can also result in differences in porosity and altered cell-ECM interactions (for example, differential presentation of RGD ligands as a function of increased ECM content). To enable more controlled studies and to test a wider range of stiffness conditions, we transitioned to the use of polyacrylamide (PA) hydrogels. Here, we synthesized PA hydrogels by varying acrylamide and bis-acrylamide concentrations, which tuned the stiffness of the hydrogels as expected (Figure 2A). Since PA is biologically inert, we covalently tethered collagen IV and fibronectin, which are crucial for adhesion of iPSC-derived BMEC-like cells^26^. We chose to synthesize PA hydrogels with stiffness conditions that represent: 1) bulk brain stiffness (0.2-2 kPa), 2) healthy peripheral vessel intimal stiffness (20-30 kPa), and 3) diseased peripheral vessel intimal stiffness (>100 kPa)^17,24^.

Here, we utilized newer variants of our iPSC differentiation protocols that replace the serum component in the culture medium with more defined B27 supplement^26^. However, we subsequently determined that iPSC-derived BMEC-like cells did not adhere to PA hydrogel surfaces as confluent monolayers unless serum (PDS) and ROCK inhibitor (Y27632) were included in the medium (Figure 2B). Serum and Y27632 have been shown to promote cell survival during passaging^38^ and were clearly necessary for attachment to the synthetic hydrogels (Figure 2C). We thus moved forward with this system for downstream analyses of cell phenotype and function. To avoid confounding effects of serum, we decreased its concentration to 0.1% and only included it for one day to assist with adherence to the PA hydrogels. Likewise, because ROCK inhibitors influence mechanosensing, Y27632 was only included for one day. All subsequent analyses were conducted 8 or 15 days after placing the BMEC-like cells onto hydrogels.

### Phenotypic and functional assessments of iPSC-derived BMEC-like cells cultured on PA hydrogels

Peripheral endothelial cell mechanobiology studies have shown that actin reorganizes in response to substrate stiffness and that this reorganization can disrupt endothelial barriers^18^. Quiescent endothelium is characterized by a cortical actin ring to reinforce cell-cell junctions, while intracellular actin stress fibers are associated with barrier dysfunction and increased permeability^2^. With this in mind, we assessed if actin organization in BMEC-like cells was affected by substrate stiffnesses of 1, 20, and 150 kPa, mimicking the conditions mentioned in the previous paragraph. After 8 and 15 days of culture on hydrogels, phalloidin staining revealed that BMEC-like cells cultured on 1 and 150 kPa substrates developed significantly higher numbers of intracellular actin stress fibers relative to cells cultured on 20 kPa substrates (Figure 3A-B). When comparing BMEC-like cells on 1 and 150 kPa substrates, the number of total stress fibers is not statistically different, but actin filaments on the stiffer matrix appear thicker and more pronounced than the diffuse filaments on the soft matrix (Figure 3A). Since complex tight junction structure is an important hallmark of the BBB, and loss of tight junction integrity and passive barrier function are typically associated with increased tight junction width and tortuosity in peripheral endothelium^12,13,33^, we quantified width and tortuosity of occludin, a major constituent of BBB tight junctions. At 3, 8, and 15 days of culture on PA hydrogel substrates, no statistical differences were observed in occludin width or tortuosity (Figure 3C-D). Next, since molecular transporters are another key aspect of BBB function, we quantified p-glycoprotein activity using fluorescent rhodamine 123 (substrate) and cyclosporin A (inhibitor) after 8 days of culture on PA hydrogel substrates, where the inhibitor yields an increase in substrate accumulation if the transporter is active. Here, we did not observe any significant difference in p-glycoprotein activity across the different substrate stiffnesses as measured by rhodamine 123 signal (Figure 3E). These results suggest passive barrier function may be compromised on softer and stiffer substrates, as judged by the formation of actin stress fibers, without overt changes to tight junction organization. In contrast, p-glycoprotein activity is unaffected by substrate stiffness.

**Figure 3.**
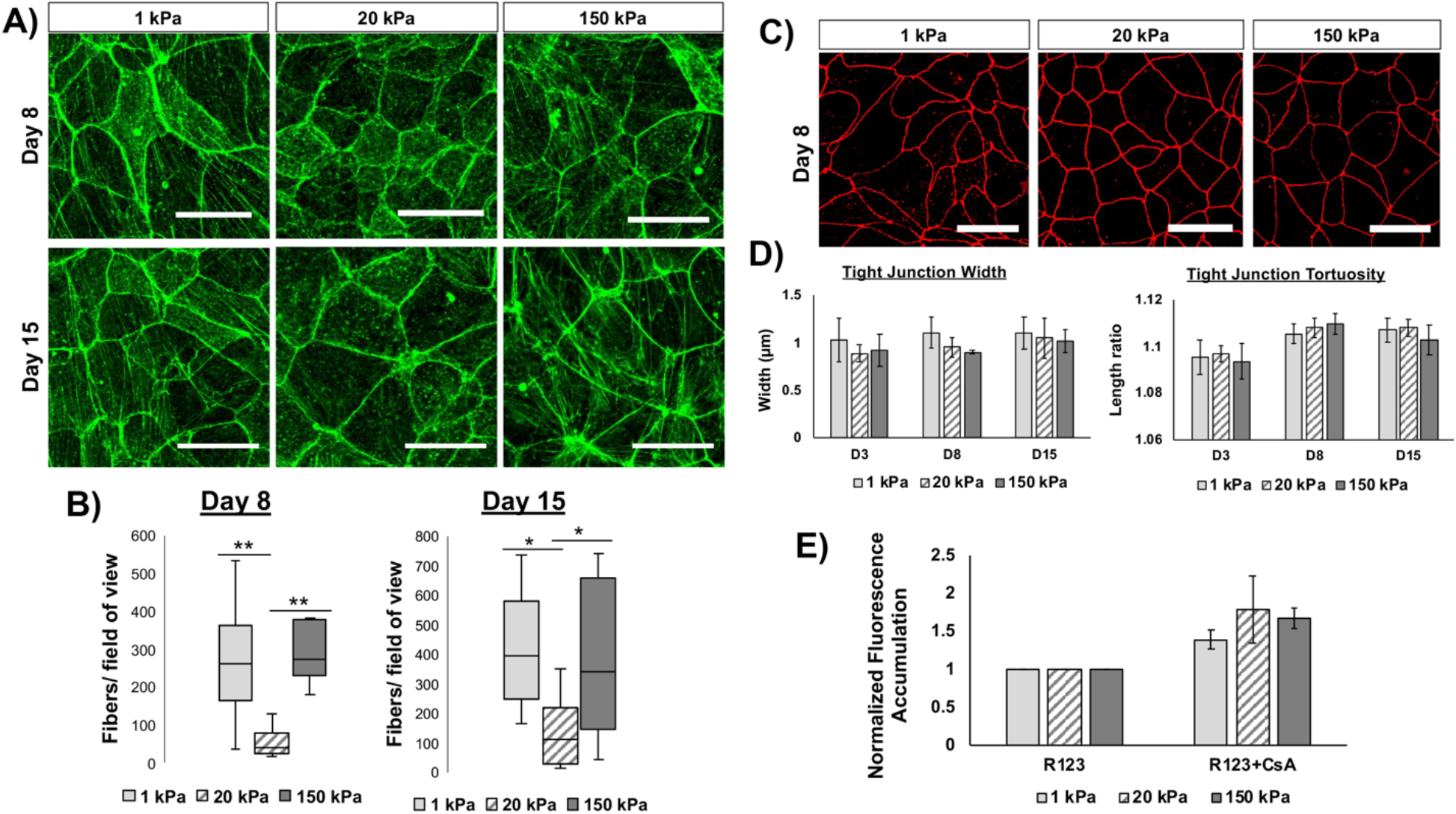
Phenotypic and functional characteristics of iPSC-derived BMEC-like cells on PA hydrogels. (A) Representative images of phalloidin-stained BMEC-like cells after 8 days and 15 days on hydrogels. Scale bars represent 30 μm. (B) Quantification of actin stress fibers on each substrate. Three distinct images were collected for each hydrogel and the total number of actin stress fibers were counted manually for each image. This process was repeated for triplicate hydrogels across 3 different iPSC differentiations (biological N=9). Data are reported as number of fibers per field of view averaged across all biological replicates. A one-way ANOVA with a Tukey post hoc test was used to evaluate the difference in actin fibers per field of view between stiffness conditions (\**p*<*0.05*, \*\**p*<*0.01*). (C) Representative images of occludin-stained BMEC-like cells after 8 days on hydrogels. Scale bars are 30 μm. (D) Quantification of tight junction width and tortuosity on each hydrogel substrate at varying time points. Three distinct images were collected for each hydrogel and tight junction width and tortuosity were analyzed using a MATLAB script. This process was repeated for triplicate hydrogels across 3 different iPSC differentiations (biological N=9). No significant differences were observed across stiffness conditions. (E) Flow cytometry quantification of rhodamine 123 (R123) uptake in BMEC-like cells after 8 days on hydrogels. Fluorescence was averaged across biological triplicates and normalized to R123 alone for each condition. Data are presented as mean ± standard deviation. No significant differences were observed in fluorescence increase upon inhibitor treatment across stiffness conditions.

### Actin disruption explicitly compromises passive barrier integrity in iPSC-derived BMEC-like cells

After observing stiffness-induced actin disruption, we assessed passive barrier function in iPSC-derived BMEC-like cells in response to a chemical modulator of actin, jasplakinolide (jasplak). Jasplak is a peptide that binds directly to actin and induces polymerization^3^ and has been shown to disrupt actin organization in endothelial monolayers^37^. Treatment of multiple endothelial cell subtypes with jasplak has been shown to diminish cortical actin signal and induce the formation of abnormal actin bundles^9^. In iPSC-derived BMEC-like cells cultured on Transwell filters without hydrogel substrates, treatment with jasplak induced a statistically significant decline in TEER in comparison to DMSO-treated and untreated controls (Figure 4A). This effect was consistent across multiple biological replicates (Figure 4B) and suggests that actin disruption can rapidly diminish passive barrier function in iPSC-derived BMEC-like cells, aligning with the actin imaging data.

**Figure 4.**
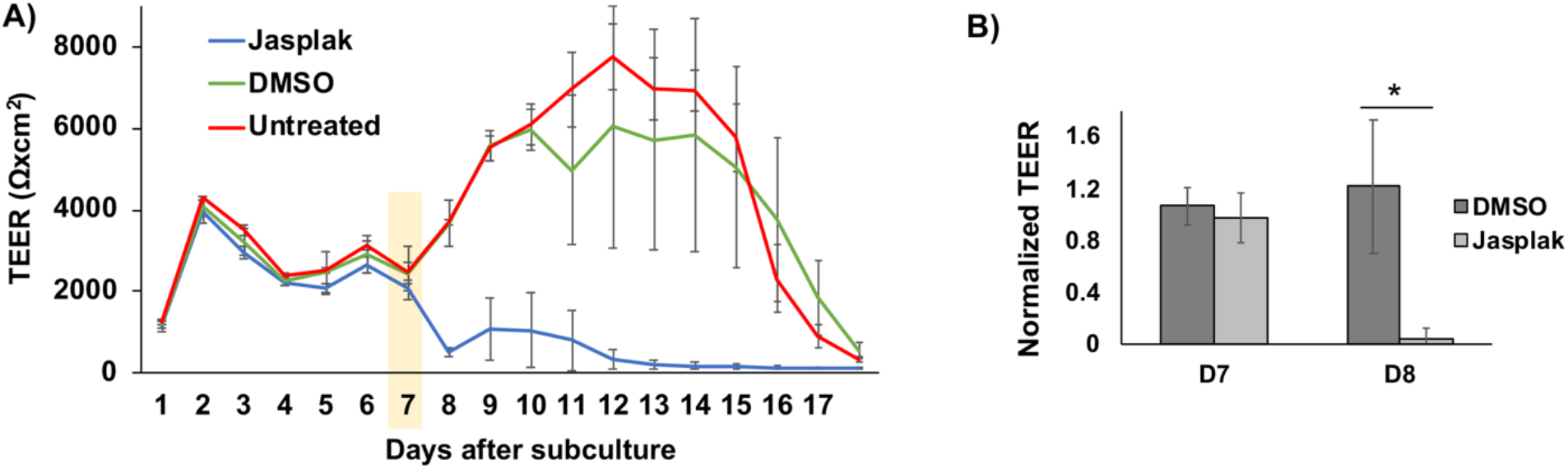
Actin disruption compromises passive barrier function in iPSC-derived BMEC-like cells independent of hydrogel substrates. (A) Representative TEER profiles in untreated cells and cells treated with jasplakinolide (jasplak) or DMSO. 3 filters were prepared for each condition, and each filter was measured in triplicate. Values for each day represent mean ± standard deviation from these 9 measurements. The yellow highlight indicates the day of treatment. (B) Average TEER before and 24 hours after jasplak or DMSO treatment. TEER values were normalized to untreated controls from the same day and averaged across 3 biological replicates (mean ± standard deviation). A student’s unpaired t-test was used to assess statistical significance (\**p*<*0.05*).

### Substrate stiffness modestly influences the transcriptome of iPSC-derived BMEC-like cells

To examine global responses to substrate stiffness, we collected RNA from iPSC-derived BMEC-like cells cultured on 1, 20, and 150 kPa substrates for 8 days. Bulk RNA-sequencing revealed a cohort of differentially expressed genes, and we used gene ontology analyses to infer differences in signaling pathway activation (Figure 5). When comparing 1 kPa to 20 kPa, 627 genes were significantly downregulated and 685 genes were significantly upregulated on the softer substrate. When comparing 20 kPa and 150 kPa, 207 genes were significantly downregulated and 218 genes were significantly upregulated on the softer substrate. When comparing 1 kPa and 150 kPa, 448 genes were significantly downregulated and 578 genes were significantly upregulated on the softer substrate. Several interesting differences were noted. In particular, cell cycle pathways are downregulated in iPSC-derived BMEC-like cells cultured on 20 kPa substrates versus 1 and 150 kPa substrates, which could reflect a more quiescent state within cells lacking actin stress fibers on a substrate stiffness mimicking healthy vessel intima. As another example, Rho GTPase signaling was upregulated on the 1 kPa substrates, and this pathway is associated with mechanotransduction and putative stress fiber formation^34^.

**Figure 5.**
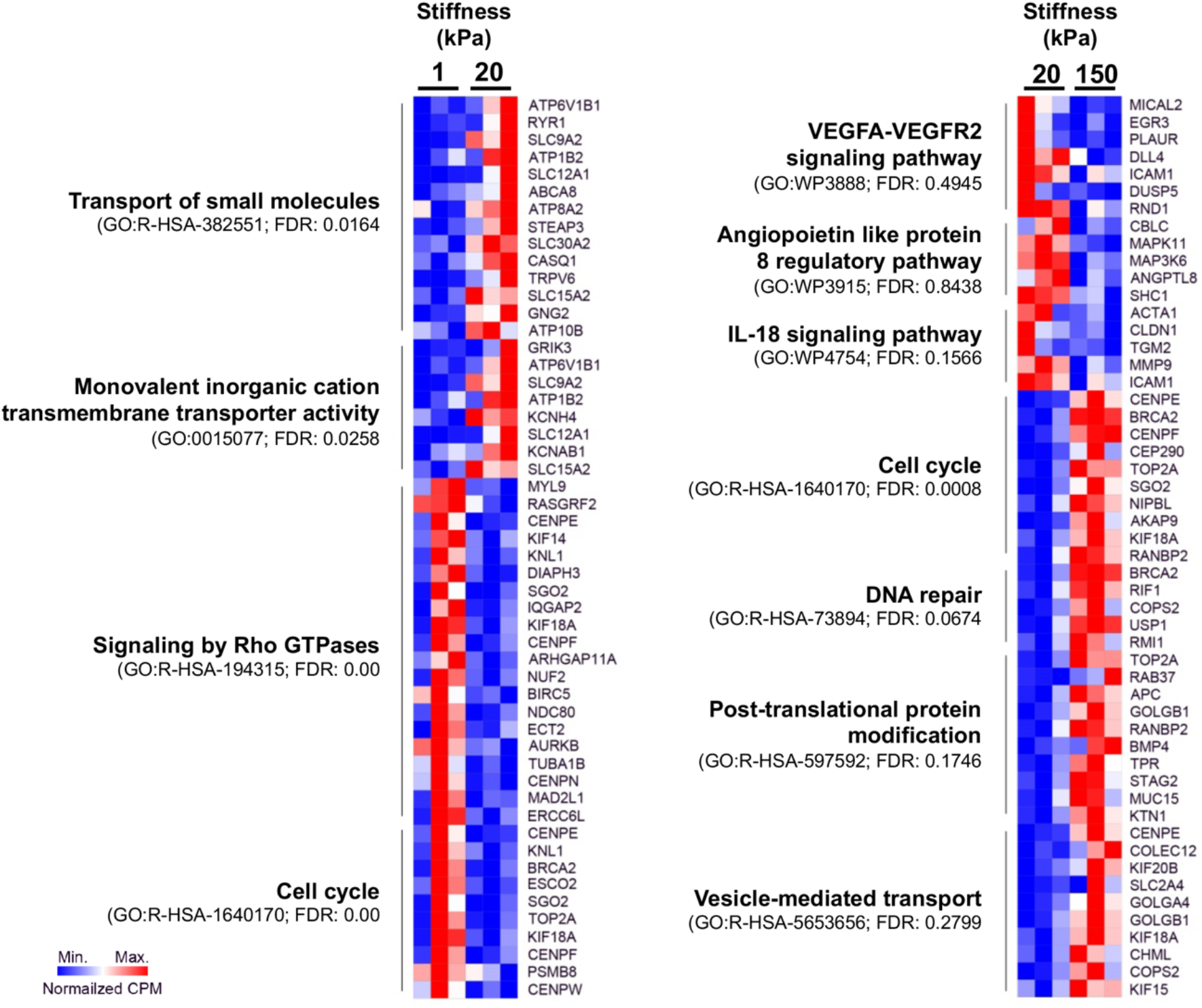
Characterization of iPSC-derived BMEC-like cell transcriptome in response to substrate stiffness. RNA was isolated and sequenced from BMEC-like cells cultured on 1, 20, and 150 kPa substrates for 8 days. Pathway enrichment analysis was performed on significantly altered genes (*p*<*0.05*). Select comparisons of 1 versus 20 kPa (left) and 20 versus 150 kPa (right) are displayed with false-discovery rates. A list of all differentially expressed genes for each pair-wise comparison are found in Table SI 1.

## CONCLUSIONS

We determined that iPSC-derived BMEC-like cells can be seeded on hydrogels for interrogation of biophysical responses to substrate stiffness. Our results demonstrate that, in agreement with general mechanobiology trends, iPSC-derived BMEC-like cells exhibit optimum passive barrier function on intermediate substrate stiffness and develop intracellular actin stress fibers on relatively soft and stiff matrices. We further demonstrate that passive barrier function is rapidly disrupted by small molecule-induced actin polymerization. Thus, similar to studies on peripheral endothelial cells^22^, endothelial cells with a highly impermeable barrier exhibit optimum function on an intermediate substrate stiffness that supports cytoskeletal health. These findings may be relevant for designing improved *in vitro* neurovascular models where BBB function must be properly represented.

Curiously, we did not observe any significant differences in tight junction morphology as a function of substrate stiffness. We have previously shown that TEER levels in iPSC-derived BMEC-like cells do not correlate with tight junction expression levels^30^, which suggests that the formation of actin stress fibers could be negatively influencing barrier function without impacting overt tight junction structures. More sensitive imaging techniques such as electron microscopy might be necessary to resolve ultrastructural differences that cannot be seen by fluorescence. We also did not observe any differences in p-glycoprotein efflux activity as a function of substrate stiffness. These results would suggest that active barrier functions are not significantly influenced by stiffness-induced cytoskeleton remodeling, but more work would be needed to probe this possibility across a more diverse array of transporters and substrates.

Overall, our results support prior investigations showing that iPSC-derived BMEC-like cells respond to mechanical cues in a manner similar to cultured endothelial cells. Our results also provide evidence that increased substrate stiffness, a ubiquitous feature of aging and disease in the periphery, may mediate negative changes in the BBB. BBB dysfunction has been linked to many disease states including stroke, neurodegenerative disease, traumatic brain injury, brain tumors, and natural aging^10,25^. In Alzheimer’s Disease (AD), BBB dysfunction is accompanied by thickening of the vascular basement membrane up to four times its original size,^23^ with increased collagen content that may alter vessel stiffness^15^. Additionally, ~90% of AD cases co-occur with cerebral amyloid angiopathy, where amyloid-β aggregates can deposit directly onto the medial layer of arteries and arterioles, disrupting vessel architecture and forming a ‘rigid cast’ around the vessel that potentially alter the mechanical properties of the vessel^16^. Hence, future work may focus on assessing whether our observations in the *in vitro* system map to human tissue samples from these patient populations.

## ACKNOWLEDGMENTS

Funding for this work was provided by a Ben Barres Early Career Acceleration Award from the Chan Zuckerberg Initiative (grant 2018-191850 to ESL), the BrightFocus Foundation (grant A20170945 to ESL), National Institutes of Health grants R21 NS106510 (to ESL), R01 NS110665 (to ESL), R61 NS112445 (to ESL), and K01 EB030039 (to KPO), and National Science Foundation grant 1846860 (to ESL). BOJ was supported by the Vanderbilt Interdisciplinary Training Program in Alzheimer’s Disease (T32 AG058524). Support for RNA sequencing was provided by the Vanderbilt VANTAGE core facility, which is supported in part by a Clinical and Translational Science Award (5UL1 RR024975), the Vanderbilt Ingram Cancer Center (P30 CA68485), the Vanderbilt Vision Center (P30 EY08126), a CTSA award from the National Center for Advancing Translational Sciences (UL1 TR002243), and the National Center for Research Resources (G20 RR030956). CTSA award UL1 TR002243 also provided pilot funding for this project. The authors would like to thank Dr. Jean-Philippe Cartailler for helpful discussions on RNA sequencing experiments, Dr. Cynthia Reinhart-King and Wenjun Wang for helpful discussions and training on polyacrylamide hydrogel synthesis, and Dr. Anthony Hmelo and John Thornton for guidance with AFM measurements.

## ETHICAL APPROVAL

No approvals were required for this study.

## DATA ACCESSIBILITY

RNA sequencing data have been uploaded to ArrayExpress under the accession number E-MTAB-10336.

